# Network-based representation learning reveals the impact of age and diet on the gut microbial and metabolomic environment of U.S. infants in a randomized controlled feeding trial

**DOI:** 10.1101/2024.11.01.621627

**Authors:** Adelle Price, Sakaiza Rasolofomanana-Rajery, Keenan Manpearl, Charles E. Robertson, Nancy F. Krebs, Daniel N. Frank, Arjun Krishnan, Audrey E. Hendricks, Minghua Tang

## Abstract

**Background:** While studies have explored differences in gut microbiome development for infant liquid diets (breastmilk, formula), little is known about the impact of complementary foods on infant gut microbiome development. Here, we investigated how different protein-rich foods (i.e., meat vs. dairy) affect fecal metagenomics and metabolomics during early complementary feeding from 5-12 months in U.S. formula-fed infants from a randomized controlled feeding trial.

**Results:** We used a novel network representation learning approach to model the time-dependent, complex interactions between microbiome features, metabolite compounds, and diet. We then used the embedded space to detect features associated with age and diet type and found the meat diet group was enriched with microbial genes encoding amino acid, nucleic acid, and carbohydrate metabolism. Compared to a more traditional differential abundance analysis, which analyzes features independently and found no significant diet associations, network node embedding represents the infant samples, microbiome features, and metabolites in a single transformed space revealing otherwise undetected associations between infant diet and the gut microbiome.

**Conclusions:** Our findings generate new hypotheses regarding the interplay between complementary feeding practices, microbial-metabolic interactions, and infant physiological outcomes. This work highlights the impact of complementary foods on infant gut microbiome development and the potential of using network representation learning to integrate multi-omic data, allowing for greater insight into complex diet, microbial, and metabolite interactions.

## Introduction

Diet has been known to have a profound impact on the gut microbiome^1–4^, which in turn plays a substantial role in host physiological development^1,5–11^. Gut microbiome establishment and development begin in infancy, when dietary differences are often attributed to whether the infant consumes breast milk or formula^12,13^. Emerging research suggests that long-term growth and immunological development, including weight gain, obesity, allergies and autoimmune diseases, can also be predicted by microbial diversity and functionality^1,5–11^. While diet-modulated differences in the microbiome of infants consuming a liquid diet (breast milk or formula) have been well-documented^12,13^, understanding of diet-modulated differences during complementary feeding (the progressive introduction of solid foods into the infant diet) is limited. The complementary feeding phase represents a critical transition period of developmental plasticity; diet choices within this time could pose both short- and long-term health consequences^14–17^. While studying the changing gut microbiome during complementary feeding holds promise, studies are challenging in part due to the need for both well controlled diet interventions, and methods that can detect more subtle microbiome changes due to diet on the background of rapid and major microbiome development often seen during this period.

To comprehensively investigate the relationship between diet, the microbiome, and infant anthropometric development, many are moving beyond traditional 16S rRNA gene sequencing to metagenomics and metabolomics. Combining metagenomics with metabolomics can enable insight into the dynamic and diverse microbiome population as well as the myriad of small, bioactive molecules produced from host metabolism of dietary components and from the microbiome through de novo synthesis^18^. While traditional ML approaches, like penalized regression, can result in overfitting and poor generalizability due to the high number of features to samples (i.e., p>>n) and high sparsity (large number of missing values where microbe and metabolite features are not observed across many samples)^19^, high-dimensional ML methods can address these issues. High-dimensional ML methods may also capture relationships between metagenomic and metabolomic features missed by traditional supervised learning ML methods.

Here, we use a high-dimensional ML method, network representation learning, to provide insight into how the gut microbiome and metabolome changes by different protein-rich foods during the introduction of complementary foods in a randomized controlled feeding trial of U.S. formula-fed infants from 5-12 months that consumed either meat or dairy-based diets. Network (or graph) representation learning has a wide range of biomedical applications including modeling protein-protein interactions, disease-gene associations, spatial organization of cells, and expanded healthcare knowledge^20–22^. Importantly, network representation learning offers several advantages for multi-omic data analysis: i) It organically integrates data types (here, metagenomics and metabolomics) by connecting them via shared samples (here, infants) and, in contrast to analyzing one feature at a time, it uses the entire multi-omics dataset to capture patterns in how all features and samples are related to each other; ii) While data sparsity is a hinderance to matrix-based operations for capturing global patterns, network-based methods can take advantage of sparse *local* patterns and create meaningful low-dimensional numerical representations (called ‘embeddings’); these embeddings naturally lend themselves to be used as inputs to ML models for downstream tasks including sample classification; and, finally, iii) By virtue of having samples and features in the same embedding space, sample classification ML models can be easily applied to classify features. Further, closeness between microbe/metabolite features within the embedding space may also point to putative functional relationships between the features. We show here that, by integrating and modeling metagenomic and metabolic data using network representation learning, we can elucidate differences in the diet groups and in turn generate new hypotheses regarding the interplay between complementary feeding practices, microbial-metabolic interactions, and infant physiological outcomes.

## Results

### Experimental Overview

In this work, we used network representation learning to examine metagenomic and metabolomic data from a seven month long randomized controlled feeding trial of 64 exclusively formula-fed 5-months-old infants recruited from the Denver, Colorado metro area. Infants were randomized to receive protein primarily from either pureed meats (n=32) or dairy foods (n=32) such as infant yogurt, cheese, and whey protein powder (**Table 1**). Fruit and vegetable intakes were not restricted, and parents were encouraged to allow the infant’s appetite to dictate their total intake^14^. Infant stool samples used here were collected at infant age 5 months and the end of the intervention, age 12 months. We used metagenomic sequencing and untargeted metabolomic profiling on the longitudinal stool collections. Microbiome features are reported with species-level taxonomic classifications as well as Cluster of Orthologous Groups (COG)^23^, KEGG Orthology (KO)^24^, and Protein Families Database (Pfam)^25^ functional classifications. Metabolites were annotated with Chemical Abstracts Service (CAS)^26^, Kyoto Encyclopedia of Genes and Genomes (KEGG)^24^, Human Metabolome Database (HMPD)^27^, and LIPID MAPS identifiers^28^. After filtering on missingness and coefficient of variation, 17,033 microbiome and 8,867 metabolome features were used for analysis (**Figure 1a**). See **Methods** for a complete description.

**Table 1.** Subject Characteristics at 5 months old^14^.

|  | Meat* (n=32) | Dairy* (n=32) | P value |
| --- | --- | --- | --- |
| Sex | 45% male | 48% male | 0.55 |
| Race and ethnicity | 75% White<br>19% Hispanic<br>3% Black<br>3% Asian | 75% White<br>16% Hispanic<br>6% Black<br>3% Asian | 0.88 |
| Birth weight (kg) | 3.31 ± 0.37** | 3.33 ± 0.48** | 0.85 |
| Gestational age (wk) | 39 ± 1 | 39 ± 1 | 0.30 |

\*Meat: meat-based complementary protein group; Dairy: dairy-based complementary protein group,
\*\*Mean ±SD; Chi-squared test used to compare proportions Independent Student's *t* test used to compare means.

**Figure 1.**
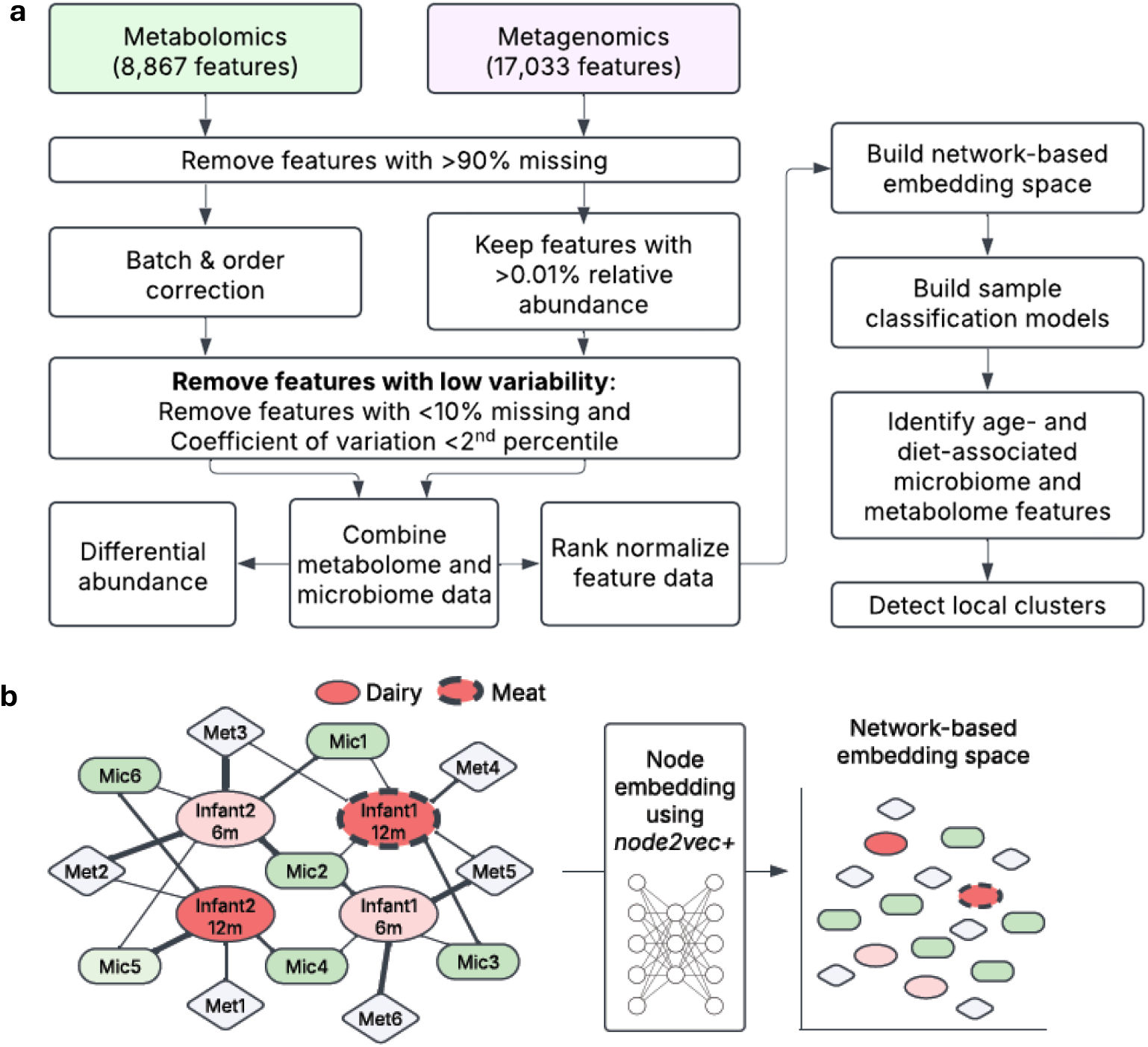
Experimental Overview to identify differences in fecal microbiome and metabolomics by age and diet. **a**, Study design and processing pipelines for microbiome and metabolite data. **b**, Schematic representation of node embedding: Input network into *PecanPy* (left), samples and features are mapped into an embedding space using weighted random walks (middle). Projection of embedding space onto a two-dimensional plane (right) shows sample-sample, feature-feature, and sample-feature relationships as spatial distances.

To identify differences in the microbiome and metabolome landscape of infants who consumed meat or dairy diets, we began with a classical differential abundance analysis examining each feature independently. We then used network representation learning^29,30^, a ML and graph theory method, to assess relationships between our sparse, high-dimensional microbiome and metabolomic data with the infant characteristics of age and diet. For the network representation framework, we performed the following steps: i) Constructed a simple network with edges connecting infant samples to microbe and metabolite features with edges weighted by the normalized abundance of the feature in the infant observation, ii) Implemented node embedding using *node2vec+ in PecanPy*^31,32^ to create a 128-dimensional numerical representation (the ‘embedding space’) of the sample-microbe-metabolite network so that the infant samples, and microbe and metabolite features were in the same transformed space; seven node embedding spaces were created using different hyperparameters to assess consistency of relationships, iii) Classified infant age or diet group using penalized logistic regression on infant node embeddings for each of the seven spaces, iv) Predicted age and diet group for each microbe and metabolite feature using the fitted infant sample classification models, and v) Performed Louvain clustering of microbe and metabolite features with consistent classification within embedding space one (**Figure 1b**).

### Differential abundance analyses identify changes in infant microbiome and metabolome by age but not between diet types

Differential abundance analysis of the log2 transformed 17,033 microbiome and 8,867 metabolite features between the 5-month-old and 12-month-old (n=50) age groups using paired Wilcoxon-rank sum tests revealed 2,493 and 1,766 significantly (FDR<0.05) higher fold changes in microbiome and metabolite features, respectively, for the 5-month-old group and 5,132 and 946 significantly higher fold changes in microbiome and metabolite compound features, respectively, for the 12-month-old group (**Figure 2a-b, Supplementary Tables 1** and **2**). Interestingly, this showed more microbiome but fewer metabolite features with a significantly higher fold change in the twelve-month group compared to the five-month group. Enrichment analysis of the significant microbiome taxa by age identified family-level enrichments in the 5-month-old age group for *f_Enterobacteriaceae* (FDR=3.0e-09). The 12-month-old age group was significantly enriched with family-level taxa *Lachnospiraceae* (FDR=1.9e-07) and *Oscillospiraceae* (also known as *Ruminococcaceae*; FDR=1.6e-05), and genera *Blautia* (FDR=3.9e-03). Numerous microbiome feature enrichments were found in both the 5- and 12-month-old age groups (**Supplementary Table 5**). Top enrichments in the 5-month-old group included genes encoding transcription factors (FDR=1.8e-12), proteins for inorganic ion transport and metabolism (FDR=1.5e-10) and *Escherichia coli* biofilm formation (FDR=1.4e-09), and transporter proteins (FDR=4.8e-09). Top enrichments in the 12-month-old group included genes encoding proteins for translation (FDR=7.4e-09), cell growth (FDR=9.8e-06), as well as ribosomal proteins (FDR=1.9e-04). All other enrichments are provided in **Supplementary Table 5**.

**Figure 2.**
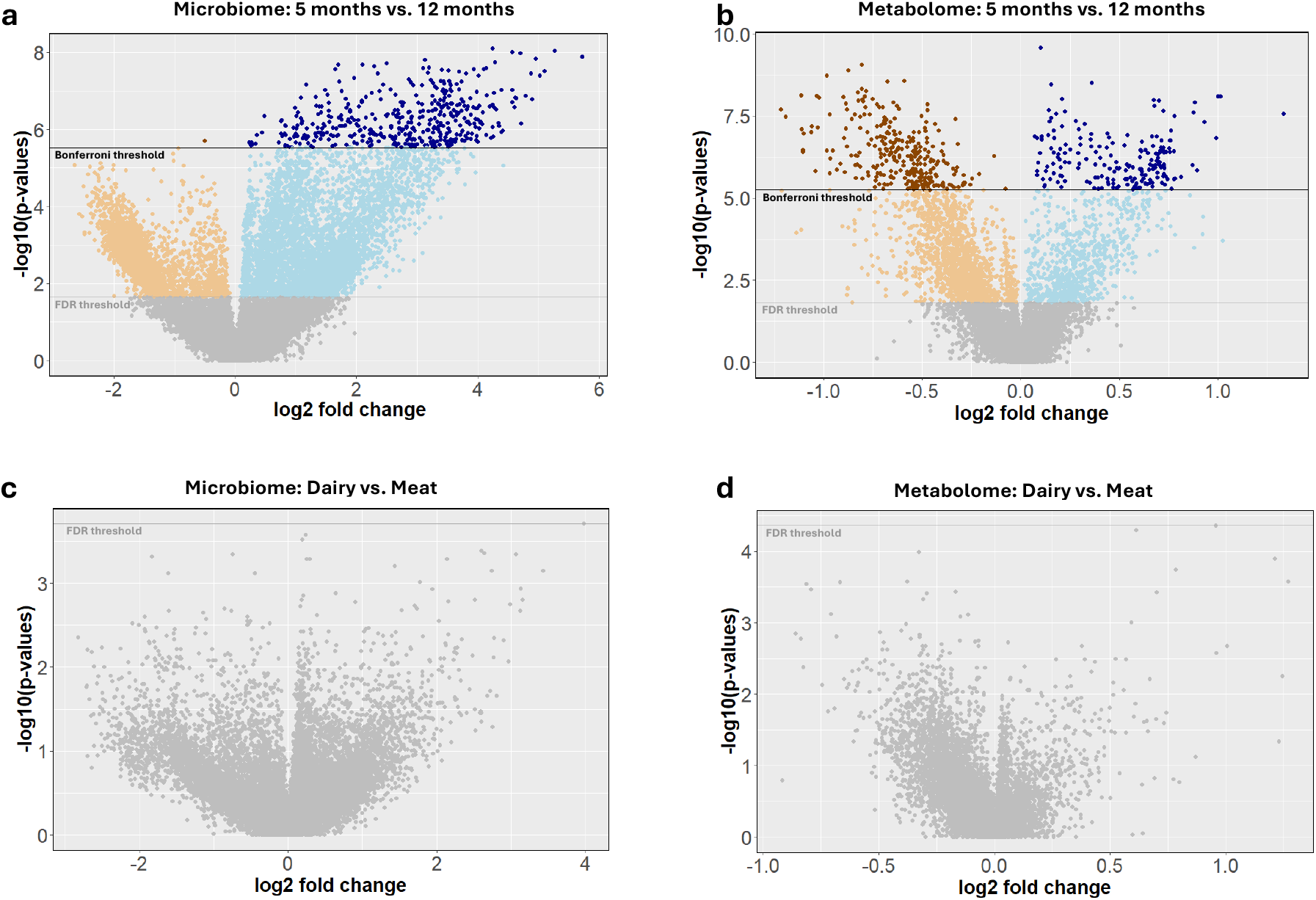
Differences in microbiome and metabolome over time and by diet. Differential abundance analysis between **a**, microbiome and **b**, metabolome features in 5-month-old and 12-month-old groups. Positive log2 fold changes indicate higher feature abundance in the 12-month-old group and negative log2 fold changes indicate higher abundance in the 5-month-old group. Differential abundance values of **c**, microbiome and **d**, metabolome features in dairy and meat groups with positive log2 fold changes indicate higher feature abundance in meat groups; negative log2 fold changes indicate higher feature abundance in dairy groups. In all figures, light orange and blue indicate passing the FDR threshold and dark points indicate passing the Bonferroni correction threshold.

We also assessed differential feature abundance of the 17,033 microbiome and 8,867 metabolite features between meat and dairy diet groups at 12 months old (*n*_*meat*_=24, *n*_*dairy*_=30 for microbiome features; *n*_*meat*_=26, *n*_*dairy*_=29 for metabolite features), using Man-Whitney U tests. We found no significant differences after controlling for multiple testing (FDR<0.05) for microbiome or metabolite features. The dairy group had 432 microbiome and 626 metabolite features with nominally significant (p_unadj_<0.05) differences compared to meat. The meat group had 553 microbiome and 169 metabolite features with nominally significant differences compared to dairy (**Figure 2c, d, Supplementary Tables 3** and **4**).

### Node embedding identifies microbiome and metabolite differences between meat and dairy diet groups in addition to age group

PERMANOVA^33^ tests of Principal Coordinates Analysis (PCoA) of the 128 node embedding dimensions by diet group yielded more distinct clustering compared to PCoA of non-embedded data, suggesting that the node embeddings typically captured slightly more variation between meat and dairy diet groups compared to the non-embedded feature data. Embedding R^2^ ranged from 0.034 to 0.062 with a median embedding R^2^ of 0.053 vs. the non-embedded R^2^ of 0.045 (**Figure 3a, b**; **Supplementary Table 6**). While we observed better performance for the age groups for both methods compared to diet groups, we observed slightly worse performance for the embedded than the non-embedded data. Age groups

**Figure 3.**
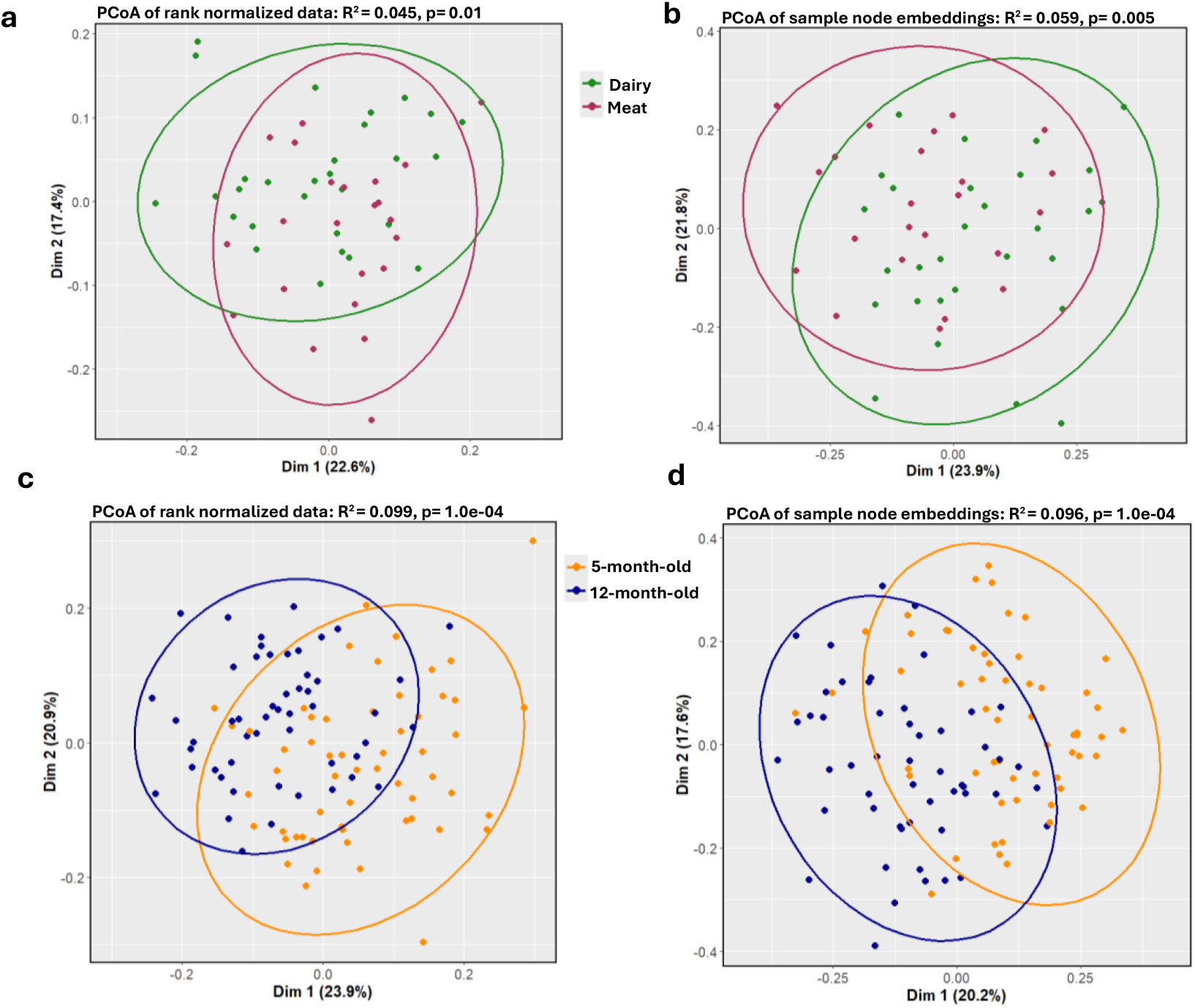
Variation between age and diet groups. PCoA of **a**, rank normalized microbiome and metabolome feature data and **b**, sample nodes in embedding space 1 across dairy (green) and meat (purple) groups. PCoA of **c**, rank normalized microbiome and metabolome feature data and **d**, sample nodes in embedding space one across 5-month-old (orange) and 12-month-old (blue) groups. embedding R^2^ ranged from 0.031 to 0.1 with a median embedding R^2^ of 0.092 vs. the non-embedded R^2^ of 0.099 (**Figure 3c, d**; **Supplementary Table 6**).

To further assess the utility of the embedding spaces, we built penalized regression models using all node embedding dimensions (n=128) to classify infant samples by age group and diet type (see **Methods**). As a point of comparison, we built a diet type and an age group penalized regression classifier using the non-embedded, rank normalized microbiome and metabolome features (n=25,900). Node embedding models had ~8% higher median F1 5-fold cross-validation (CV) validation score for diet group sample classification compared to rank normalized models (median F1 CV fold validation score=81% and 73%, respectively). Conversely, node-embeddings had similar, although slightly lower, performance compared to non-embedded models for discriminating age group (median F1 CV fold validation score= 89% and 91%, respectively). Notably and as expected from the PCoA results, model performance was again much higher for age group classification compared to diet type classification across all models. This suggests that the non-embedded rank normalized data may be sufficient to distinguish larger effects in the microbiome and metabolome space (such as those due to age) while the node embedding space may enable detection of more complex or subtle differences, such as those due to diet.

### Node embedding identifies diet-associated microbiome and metabolome features

Beyond infant sample classification, the low-dimensional node embedding space that contains both infant samples and microbiome and metabolome features presents a crucial benefit: microbiome and metabolome features can be assigned to a most likely diet and age group based on their location in the node embedding space. To account for potential variability in embedding space generation, we assigned features to age and diet groups if the classification model predictions using a 0.5 probability threshold were consistent across all seven node-embedding spaces.

Out of 8,867 metabolite and 17,033 microbial features, 206 metabolite and 785 microbial features were consistently classified into the meat group, and 21 metabolite and 85 microbial features were classified into the dairy group (**Supplementary Table 7**). Hypergeometric enrichment analysis of features classified into the meat group revealed COG enrichments of translation, ribosomal structure and biogenesis (FDR= 2.43e-08), nucleotide transport and metabolism (FDR=0.005), amino acid transport and metabolism (FDR=0.006), carbohydrate transport and metabolism (FDR=0.02), and DNA replication, recombination, and repair (FDR=0.03). Among Pfam features, the meat diet group was enriched at nominal significance in protein families performing DNA repair (FDR=0.07, p_unadj_=0.02), translation (FDR=0.07, p_unadj_=0.03), and ribosomal proteins (FDR=0.07, p_unadj_=0.01). Additionally, the meat diet group was enriched at nominal significance in KO genes involved in histidine metabolism (FDR=0.07, p_unadj_=0.004). Notable metabolites classified into the meat group that have been previously shown to be associated with diets high in animals fats or protein including Taurocholic Acid, a bile acid with taurine and cholic acid components^34,35^, 4-Hydroxyphenylpyruvic Acid (4-HPPA), an intermediary compound in tyrosine metabolism^36,37^, Leukotriene F4, a derivative of arachidonic acid (a fatty acid abundant in meat)^38,39^, and Deoxyinosine, a purine derivative^40,41^ (**Supplementary Tables 7** and **8**). No enrichments were found in the dairy group. Notable metabolites classified into the dairy group included D-erythro-Sphingosine C-20, a sphingoid base derived from sphingolipids, which are abundant in dairy products^42^, and C16-OH Sulfatide, a glycosphingolipid also present in dairy products^42^ (**Supplementary Tables 7** and **8**).

Within the age groups, 387 metabolite and 798 microbial features (including 7 microbial species) were classified into the 5-month-old group, and 199 metabolite and 926 microbial features (including 47 microbial species) were classified into 12 months (**Supplementary Table 10**). Enrichment analysis of features at 5 months showed significant enrichments of KO genes encoding transcription factors (FDR=0.002), biosynthesis of siderophore group nonribosomal peptides (FDR=0.003), and benzoate degradation (FDR=0.04) (**Supplementary Table 11**). Notably, some metabolites classified into the 5-month-old group have been previously shown to decrease in the first year of life including Testosterone sulfate, a form of the hormone testosterone^43^, and Cortisol 21-sulfate, a form of the hormone cortisol^44^ (**Supplementary Table 10**). At 12 months, significant enrichments were found in COG features for cell motility (FDR=0.005) and defense mechanisms (FDR=0.02), KO features encoding proteins involved in cell growth (FDR=0.006) and amino sugar and nucleotide sugar metabolism (FDR=0.04). Additionally, the 12-month-old group was enriched in microbial species within the family *Oscillospiraceae* (FDR=2e-04) and class *Clostridia* (FDR=1.01e-05) (**Supplementary Table 11**).

### Microbiome and metabolite features form diffuse clusters by function in node embedding space

Just as the spatial landscape of the node embeddings allows classification of microbiome and metabolome features to infant characteristics of age and diet groups, the relative closeness between features within the embedding space may also be biologically meaningful. Embeddings of two features (microbe-microbe, metabolite-metabolite, or microbe-metabolite) are close to each other when they have highly similar patterns of normalized abundances across infant samples with the same characteristics (e.g. across all infants with a meat diet). To explore spatial relationships within the embedding space, we created a feature similarity network with the edge weight between two features equal to the distance of their embeddings. We then performed Louvain clustering on the spatial network and identified functional enrichments within each cluster for diet and age groups within embedding space one (**Supplementary Table 6**).

Across all time and diet groups, Louvain clustering identified three clusters that were highly interconnected and not strongly defined. Modularity of the clusters (describing how well a network is partitioned into groups according to within-cluster connection density and between-cluster connection sparsity; typically ranging from 0 to 1^45^), ranged from 0.087 within features classified into the 12-month-old group to 0.161 within the 5-month-old group (modularity= 0.128 and 0.113 for features within the dairy and meat groups, respectively). Notably, though clusters were not strongly defined, functional enrichment analysis by Louvain cluster showed more within-cluster enrichments compared to enrichment analysis across all features classified into a time or diet group indicating microbiome and metabolite features tended to group by function within the embedding space.

For the 206 metabolite and 785 microbial features classified into the meat group, the largest cluster within the meat group was driven primarily by microbiome features (479 microbiome, 61%, vs. 43 metabolite, 21%) involved amino acid metabolism (histidine, cysteine and methionine pathways enriched at FDR<0.05), translation and ribosomal proteins, and DNA metabolism (enrichments in replication, recombination, repair, and nucleotide metabolism). The second cluster contained the largest proportion of metabolite compounds with 143 metabolites (69%) and 261 microbiome (33%) features and was functionally enriched in proteins for transport, secretion systems, cell processes, and cell wall/membrane/envelope biogenesis, as well as DNA binding and integration functions. The third and smallest cluster with 20 metabolite (10%) and 45 microbiome (6%) features was enriched in carbohydrate transport and metabolism (**Figure 4a, Supplementary Table 11**). For the 21 metabolite and 85 microbial features classified into the dairy group, only one Louvain cluster with 3 metabolite and 36 microbiome features was functionally enriched for transporter proteins (**Supplementary Table 12**).

**Figure 4.**
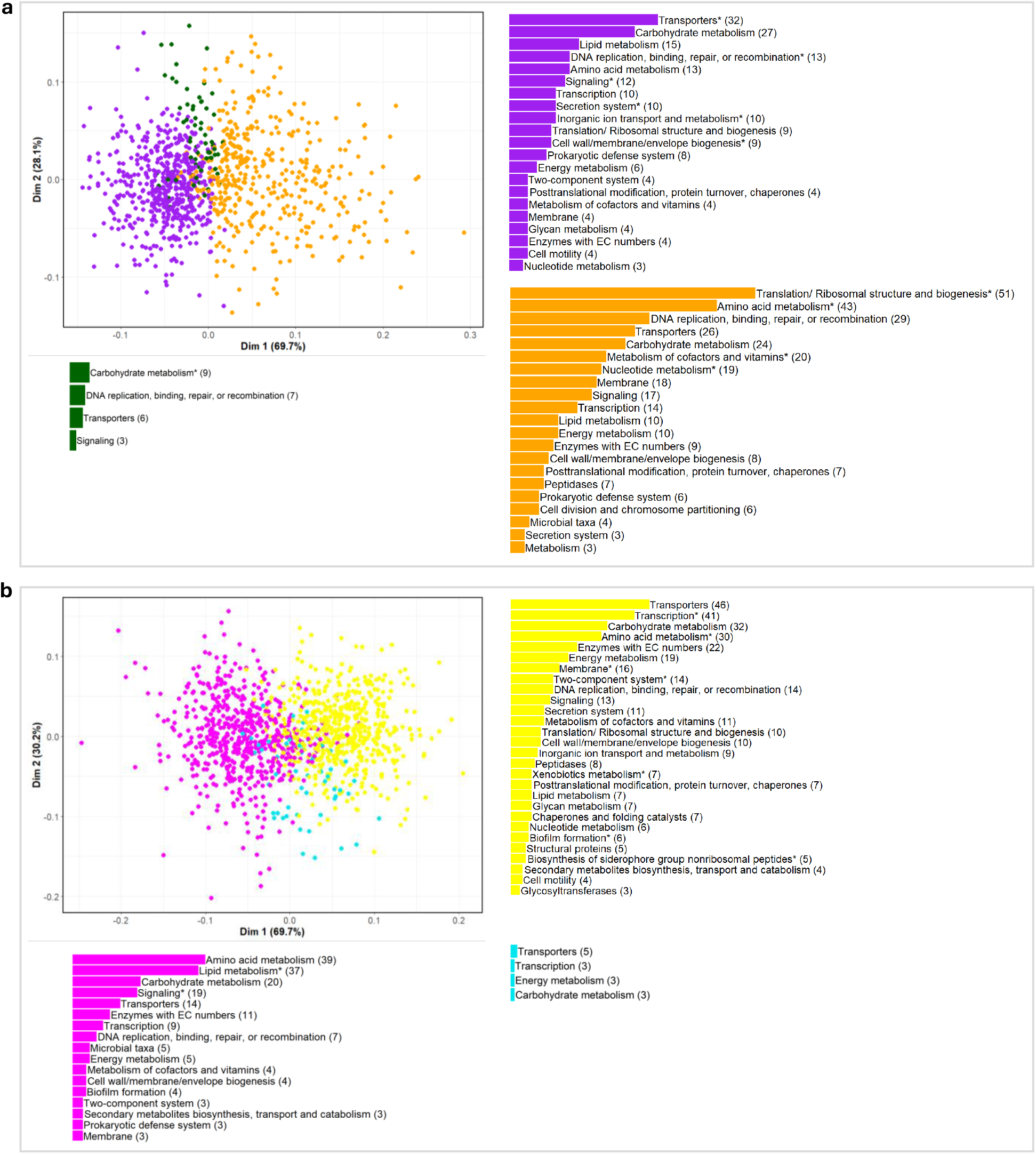
Putative functional clusters of microbiome features and metabolites. PCoA of microbiome and metabolite feature embedding vector pairwise cosine similarity within embedding space one for features classified into the **a**, meat and **b**, 5-month-old groups with feature count per broad functional category, colored by cluster (see **Supplementary Table 15**). Broad functional categories with at least one FDR significant functional enrichment noted with an asterisk.

For the 5-month-old group with 387 metabolite and 798 microbiome features, the largest cluster was again driven by microbiome features with 528 microbes (66%) and 27 metabolites (7%) and contained 14 enrichments including in transcription factors, peptides for biosynthesis of siderophore groups, and membrane proteins. The second cluster contained 357 metabolite (92%) and 171 microbiome (21%) features and was enriched in genes encoding proteins for quorum sensing and steroid hormone biosynthesis (**Figure 4b, Supplementary Table 13**). For the 199 metabolite and 926 microbiome features classified into the 12-month-old group, the first cluster contained a relatively high proportion of metabolites (81%, 162) and approximately the expected third of microbial features (27%, 251) with enrichments in sphingolipid and nucleotide sugar metabolism as well as signal transduction mechanisms. The second cluster with 26 metabolite (13%) and 374 microbial (40%) features was enriched in family-level taxa *f_Oscollospiraceae* as well as genes encoding proteins for energy, carbon, and methane metabolism, cell motility, and cellular defense mechanisms (**Supplementary Table 14**).

## Discussion

Here, we investigated differences in the microbiome and metabolome of infants by age and meat- or dairy-based complementary diet group. While traditional differential abundance analyses of individual microbiome and metabolite features identified features that significantly differed by age, no significant differences by diet group were identified after adjusting for multiple comparisons. This was perhaps not surprising given the relatively small sample size (*n*_*meat*_=24, *n*_*dairy*_=30 for microbiome features; *n*_*meat*_=26, *n*_*dairy*_=29 for metabolite features) and high sparsity of the microbiome and metabolome data. To improve identification of more subtle differences in the microbiome and metabolome due to different dietary interventions, and better use the sparse data, we applied network representation learning with *PecanPy*^*31*^ to model infant fecal sample-microbiome-metabolite interactions in the same embedded space. Using the embedded space, we observed improved prediction of infant sample classification into diet groups and identified numerous biological pathways enriched within diet or age groups.

In the present study, we identified enrichment of several biological pathways within the meat diet group, including microbial genes involved in amino acid metabolism. Ten percent or more of ingested protein can reach the colon, and the amount is at least partially dependent on protein quality^46^. Different protein quality between meat and dairy may result in differing availability of dietary proteins to gut microbiota. Amino acid compositions for various protein sources also differ. For example, dairy has a higher amount of branched-chain amino acids than meat^47^. Additionally, upregulation of amino acid metabolism pathways has been previously linked to omnivorous diets compared to vegan and vegetarian diets in adults^48^. This different amino acid metabolism by the gut microbiota in meat vs. dairy groups may be related to the previously observed growth differences between the two groups^14^. Moreover, gut microbial amino acid metabolism is suspected to play a role in the development of obesity and Type 2 Diabetes Mellitus in adults^49–51^, suggesting a potential role in health; although more research is needed to fully investigate the relationship between diet, amino acid metabolism and health overall.

We also explored microbial and metabolite feature cluster formation within our embedding spaces to identify potential functional relationships within the contexts of diet and age. Clustering analysis on the spatial distance between microbial features and metabolites within the embedding space revealed features tended to cluster by function. For all time and diet groups, we found more functional enrichments within specific Louvain clusters compared to enrichment using all features classified into a given time or diet group. Notably, within features classified into the meat diet group, we found further evidence for enrichments of amino acid metabolism pathways including those of histidine, cysteine, and methionine metabolism. Future mechanistic research is needed to identify the linkage between health indicators and these amino acid metabolism pathways, which may serve as a mediator between diet and infant health outcomes; possibly offering an opportunity for early intervention using gut microbiota as a therapeutic target^15–17^.

For the age group comparisons, we found traditional differential abundance analyses identified many microbial and metabolic features differing by age, confirming that changes in microbiome in infants over ages 5 to 12 months are large and readily detectable^52,53^. The microbiome of the 5-month-old infants was characterized by enrichment of species in the *Enterobacteriaceae* family. This is consistent with prior studies showing *Enterobacteriaceae* is among the earliest colonizers of the infant gut^54,55^. At 12-months-old, we found the infant gut microbiome was enriched for family-level taxa including *Lachnospiraceae* and *Oscillospiraceae* (*Ruminococcaceae*), which have been shown to increase greatly in first year of life^53,56–58^. Features classified into the 12-month-old group across the embedding spaces were also enriched in family-level taxa *Oscillospiraceae* (*Ruminococcaceae*).

Studies examining the effects of complementary foods on gut microbiome, especially for the full spectrum of microbial taxa, to genetic features, to fecal metabolites, are limited. Here, we examined gut microbiome and fecal metabolite changes in response to a randomized controlled feeding trial of meat and dairy-based complementary diet intervention in U.S. infants from 5 to 12 months. We applied node embedding with network representation learning to the high-dimensional, complex landscape of microbes and metabolites allowing for a deeper exploration of interactions between microbes, their metabolites and infant protein source and age, enhancing our understanding of diet and the gut microbiome during infant development.

## Methods

### Study design and human subjects

Stool samples from a completed randomized controlled feeding trial^14^ were used in this study. In brief, formula-fed infants (n=64) were recruited from the metro Denver area using mail-in flyers by the Colorado Department of Public Health. After informed consent was obtained from participants’ legal guardians, participants were randomized to consume meat (n=32) or dairy (n=32) as the primary source of protein from the complementary foods. Exclusion criteria included low birth weight, cumulative breastfeeding > 1 month, and significant congenital anomalies or known chronic diseases. The primary growth outcome was presently previously^14^. The randomized controlled feeding trial was conducted according to the guidelines of the Declaration of Helsinki, approved by the Colorado Multiple Institutional Review Board, and registered at ClinicalTrials.gov (NCT02142647; Study registration date: 2014-05-14).

After stratifying by sex and ethnicity, 5-month-old, exclusively formula-fed infants were randomized to receive either pureed meats or dairy foods such as infant yogurt, cheese, and whey protein powder. Participants visited the Pediatric Clinical & Translational Research Center (CTRC) at Children’s Hospital Colorado (CHCO) at baseline (5 months) and at the end of the intervention (12 months). Compliance and health history were obtained at the monthly home visits. Stool samples were collected at 5, 9 and 12 months.

Metabolome features were isolated from 5-old-months (n=60), 9-months-old (n=52), and 12-months-old infant samples (*n*_*meat*_=26, *n*_*dairy*_= 29), with 167 infant samples total. Microbiome features were isolated from 5-old-months (n=57) and 12-months-old infant samples (*n*_*meat*_=28, *n*_*dairy*_= 29), with 122 infant samples total. See below for more information.

### Dietary intakes

During the 5 to 12 months dietary intervention, infants consumed a standard, intact milk-protein-based formula *ad lib*. Infant formula (Similac Optigro, Abbott Nutrition, IL) was provided during the intervention to standardize this exposure. The meat-based diet consisted of commercially available pureed meats, and the dairy-based diet consisted of infant yogurt, cheese, and a powdered concentrate of 80% whey protein (specially packaged for this study by Leprino Foods (Denver, CO). Fruit and vegetable intakes were not restricted. Parents were provided tailored feeding guidelines and were encouraged to let the infant’s appetite dictate their total intake, as done with previous complementary feeding interventions by our group^59–61^. Three 3-day diet records (de-identified) at 5, 9 and 12 months were analyzed by the CTRC Nutrition Core (NDSR software, Minneapolis, MN).

### Stool sample collection

To collect stool samples, sterile diaper liners, gloves and pre-labeled ziplock bags were given to caregivers to place in the infant’s diaper. The liner was biodegradable, which effectively collected stool, but allowed passage of urine. Caregivers were given instructions on how to place and remove the liner from the diaper once stool was produced. The soiled liner was then placed by the caregiver in ziplock bags with written collection time and date. The caregiver placed the bagged samples in home freezers (−20° C) and notified the study coordinator immediately. The study coordinator collected the samples within 24 hours and transferred them to University of Colorado Anschutz Medical Campus on ice packs. Stool samples were then collected by research personnel from the liner in a laminar flow hood and stored in sterile vials in −80° C freezers until analysis.

### Fecal untargeted metabolomics

Fecal homogenates (50ul) were deproteinzied with six times volume of cold acetonitrile:methanol (1:1 ratio), kept on ice with intermittent vortexing for 30 minutes at 4^°^C, then centrifuged at 18000xg. 13C6-phenylalanine (3 µl at 250ng/µl) was added as internal standard to fecal homogenates prior to deproteinization. The supernatants were divided into 4 aliquots and dried down using a stream of nitrogen gas for analysis on a Quadrupole Time-of-Flight Mass Spectrometer (Agilent Technologies 6550 Q-TOF) coupled with an Ultra High Pressure Liquid Chromatograph (1290 Infinity UHPLC Agilent Technologies). Profiling data was acquired under both positive and negative electrospray ionization conditions over a mass range m/z of 100 - 1700 at a resolution of 10,000 (separate runs) in scan mode. Metabolite separation was achieved using two columns of differing polarity, a hydrophilic interaction column (HILIC, ethylene-bridged hybrid 2.1 × 150 mm, 1.7 mm; Waters) and a reversed-phase C18 column (high-strength silica 2.1 × 150 mm, 1.8 mm; Waters) with gradient described previously^62–64^. Run time was 18 min for HILIC and 27 min for C18 column using a flow rate of 400µl/min. A total of four runs per sample were performed to give maximum coverage of metabolites. Samples were injected in duplicate, wherever necessary, and a pooled quality control (pooled QC) sample, made up of all of the samples from each batch, was injected several times during a run. A separate plasma quality control (QC) sample was analyzed with pooled QC to account for analytical and instrumental variability. Dried samples were stored at −20^°^C till analysis. Samples were reconstituted in running buffer and analyzed within 48 hours of reconstitution. Auto-MS/MS data was also acquired with pooled QC sample to aid in unknown compound identification using fragmentation pattern.

Data alignment, filtering, univariate, multivariate statistical and differential analysis was performed using Mass Profiler Professional (Agilent Inc, USA). Metabolites detected in at least ≥80% of one of two groups were selected for differential expression analyses. Metabolite peak intensities and differential regulation of metabolites between groups were determined as described previously^62,64^. Each sample was normalized to the internal standard and log 2 transformed. Default settings were used except for signal-to-noise ratio threshold (3), mass limit (0.0025 units), and time limit (9 s). Putative identification of each metabolite was done based on accurate mass (m/z) against METLIN database^65^ using a detection window of ≤7 ppm. The putatively identified metabolites were annotated as Chemical Abstracts Service (CAS)^26^, Kyoto Encyclopedia of Genes and Genomes (KEGG)^24^, Human Metabolome Database (HMBD)^27^, and LIPID MAPS identifiers^28^.

### Fecal metagenomics

DNA was extracted from 25-50 mg of each stool sample using the QIAamp PowerFecal DNA kit (QIAGEN, Inc), which employs both mechanical and chemical lysis. DNA concentrations were measured using a Qubit® 2.0 Fluorometer (Invitrogen, Inc.) following the manufacturer’s protocol for the dsDNA HS Assay Kit. The NexteraXT DNA sample preparation kit (Illumina, Inc) was used to prepare 1 ng of DNA for multiplexed shotgun metagenomic sequencing, following the manufacturer’s protocol. Paired-end 2×150 nucleotide sequencing of pooled libraries was conducted on the Illumina NextSeq 550 platform with the NextSeq High Output Kit v2.5 (Illumina, Inc). Sequences were demultiplexed by library and putative human reads culled by bowtie2^66^ mapping to the hg19 human reference genome. All resulting sequences that did not map to the human genome were deposited in the NCBI Sequence Read Archive under project accession number PRJNA884688. These non-human reads were then processed by Trimmomatic (v0.39^67^) to remove low quality bases (parameters: LEADING:3 TRAILING:3 SLIDINGWINDOW:4:15 MINLEN:50) and adapter sequences. This process produced a median of 4.8×10^6^ reads per library (IQR: 3.1×10^6^ – 6.8×10^6^ reads/library).

Quality-filtered reads were then annotated using the SqueezeMeta pipeline (v1.4.0^68^), by first co-assembling sequences across all libraries with MEGAHIT (v1.0^69^), removing short contigs (<200bp) with prinseq, and then predicting open reading frames (ORFs) in remaining contigs with Prodigal^70^. ORFs were annotated using DIAMOND^71^ to query the NCBI GenBank non-redundant database^72^ for taxonomic classification and EggNOG^73^ and KEGG databases^24^ for functional annotations of Clusters of Orthologous Groups (COG)^23^ and KEGG orthology (KO)^24^ classifications, respectively. Assignments to the Protein Families Database (Pfam)^25^ were made by Hidden Markov Model (HMM) search using HMMER3^22^.

Bowtie2^10^ was used to map reads assignments were made by Hidden Markov Model (HMM) search using HMMER3^74^. Bowtie2 was used to map reads to contigs. The final outputs of SqueezeMeta were separate, raw read count files for species-level taxonomic classifications, as well as COG, KO, and Pfam functional classifications.

### Filtering and preparation of the metabolome and metagenome data

Beginning with 37,194 microbiome features (22,656 COG; 5,733 KO; 7,109 Pfam; 1,696 Bacterial species-level taxa) across 167 infant samples, we removed observations with <250k reads or duplicates. We then removed features with more than 90% missing across all observations and maximum relative abundance among subjects <0.01%. Next, to focus on variable microbiome features we removed compounds with low variation and missingness (Coefficient of Variation (CV) < 2^nd^ percentile and missingness <10%). We retained 17,033 microbiome features for analysis.

Beginning with 9,046 metabolite features (1,556 negative aqueous; 2,409 negative lipid; 1,784 positive aqueous; 3,297 positive lipid) across 112 infant samples, we removed compounds with more than 90% missing across all observations. To correct for batch and drift effects, we used a linear regression with each compound’s abundance as the outcome and processing batch, order of sample processing within each batch, and batch by order interaction terms as predictors. Outliers as defined by the linear regression were removed and the updated fitted model was used to estimate residuals for all observations including outliers. The residuals were then shifted by the difference between the original and residual minimum value of the residuals to preserve the minimum value from the starting dataset resulting in the final batch- and order-adjusted values for each compound. To focus on variable compounds, we removed compounds with low variation and missingness (Coefficient of Variation (CV) < 2^nd^ percentile and missingness <10%). We retained 8,867 metabolite features for analysis.

We merged metabolites and microbiome feature data by infant sample time and diet label. After the merge, we retained 57 infant samples at 5-months-old and 52 infant samples at 12-months-old (with *n*_*meat*_=23, *n*_*dairy*_=29 at 12-months-old).

### Differential abundance analysis

Statistical analysis of differential abundance for the log2 transformed microbiome and metabolomic compound relative abundance was performed using paired Wilcoxon signed-rank test or non-parametric Mann-Whitney U-test for the paired age group and diet type data, respectively. Missing values (coded as 0’s) in both log2 transformed microbiome and metabolomic compounds were thought to be below the minimum detectable limit and were thus changed to the minimum value observed. An FDR correction was used to adjust for multiple testing.

### Building the network embedding

To transform the filtered microbiome and metabolome data into a weighted, undirected network as input for *PecanyPy*^31^, we rank normalized the percent relative abundance (microbiome) or compound abundance (metabolites) values across observations. We merged the microbiome and metabolite data by observational unit (infant sample), retaining 109 observations, then reshaped the data into a sample-feature pair edge list and the corresponding rank-normalized feature value as the edge weight. Note that missing values (coded as 0’s) observed in percent relative abundance or compound abundance values were kept as zeros in the ranking step to preserve the original data structure in the network (Pipeline figure in **Figure 1a**).

Next, 128-dimension network embeddings were generated using the *PecanPy*^31^ implementation within *Python* 3.10 of the *Node2Vec+* algorithm^32^. To optimize the embedding space, we used *wandb* (wandb.ai) hyperparameter sweeps to tune the *Node2Vec+* return parameter (p), in-out parameter (q), and gamma value (g). For all other parameters, the default values were used.

### Assessing infant sample separation by time and type

To assess infant sample separation within the rank normalized data or embedding spaces, we calculated a distance matrix as 1 minus the cosine similarity of microbiome and metabolome features and the infant samples. We then performed PERMANOVA^33^ analysis with 10,000 permutations using the *adonis* function within the *vegan* R package^75^.

### Building sample classifiers to evaluate embedding spaces

We trained penalized logistic regression models to classify infant samples by either age or diet group. (Diet group was only assessed at 12-months.) We designed 5 training/testing splits that were balanced for both age and diet type infant samples and trained both an age and diet classifier for each split. For each model, CV within the training split was used to select the best hyperparameters: penalty type (l1, l2 or elastic net), strength, and ratio (if penalty was elastic net). Since elastic net models did not always converge, we opted to allow all three penalty types. Next, we used the training split for each fold to train one model and evaluate the performance on the held-out test split. As an evaluation metric, we calculated one score for each embedding space: the mean F1 score on the test samples for all classifiers (5 diet-type, and 5 age group classifiers). We conducted an initial hyperparameter sweep using a range of 0.01-50 for p, 0.1-25 for q and 0,1, or 2 for g using only age group models to score embedding spaces.

Based on the results of the first sweep, we conducted a second sweep with 200 models using a range of 0.1-25 for p, 0.1-10 for q and either 0, 1, or 2 for g using both age group and diet-type classifiers. We then assessed the variance of all unique embedding spaces generated across both sweeps, 296 in total. For this, we used a combination of median F1 score and IQR for the five models of each classifier type. This resulted in seven embedding spaces with a minimum median F1 score of 0.85 and a maximum IQR of 0.25. Embedding space hyperparameters are noted in **Supplementary Table 6**.

### Building feature classifiers

For each of our seven embedding spaces, we used the original 5 folds to perform CV and select the best hyperparameters to train a final deployment model of each classifier type. Since the deployment models are used to prioritize features, we elected to use all available infant samples for training or validation rather than create a final held-out test set. We report the validation performance on each of the 5 folds as well as model hyperparameters in **Supplementary Table 6**; but note that the embedding scores should be used to assess the model’s generalizability to unseen infant samples as we did not use an unseen test set to assess the models described here (all samples were used in training, validation, or hyper-parameter tuning). As a baseline, we also trained classifiers using the same approach but with the rank normalized metabolome and microbiome features. All model training was done using *scikit-learn*^76^.

### Identification of features associated with diet and age using node embedding

The class (age class: 5-month-old or 12-month-old; or diet class: meat or dairy) of each microbiome and metabolome feature was predicted using each of the seven prediction models built on the node embeddings developed with infant samples, microbiome features, and metabolome features. Feature class was recorded as the intersection of all prediction model classes (e.g. if a feature was classified as ‘Meat’ across all seven prediction models, it is noted here as a meat-associated feature). Classification was determined using a 0.5 threshold. Average class prediction probability was calculated as the average class prediction probability across all seven prediction models.

### Clustering microbiome and metabolite features in node embedding space

To assess spatial clustering of the microbiome and metabolome features classified into diet groups within the embedding space, the distance between features within one embedding space was calculated using pairwise cosine similarity across the embedding dimensions used in the final diet or age group classifier. To focus on spatially similar features in the embedding space for clustering, we set paired feature distances >50^th^ percentile to 0, and transformed the distances with a squared term. We then performed Louvain clustering^77^ on edge weights (with edge weights set equal to pairwise distance) between all feature pairs among the features classified. We used Gephi^78,79^ open-source software to perform Louvain clustering with default parameters.

### Enrichment Analysis

Each microbial feature was annotated with functional categories using species-level taxonomic, as well as COG, KO, and Pfam functional classifications collected during Fecal Metagenomics data extraction (see **Methods**). Pfam functional classifications were obtained through the Gene Ontology^80^ mapping of GO terms to Pfam entries (https://current.geneontology.org/ontology/external2go/pfam2go; version date 2024/01/31, 08:33:46). KEGG functional pathways were obtained through the KEGG Mapper search tool^81,82^ mapping of KEGG annotated microbiome features (KO) and KEGG annotated metabolites (https://www.kegg.jp/kegg/mapper/search.html; last updated: July 12, 2021). Among features and within each cluster, we filtered to annotated functional categories with at least four features. We then created a contingency table (containing the number of features in the functional category of interest within the target group, ie. Meat, the number of features not in the functional category of interest within the target group, number of features in the functional category of interest within the alternative group, e.g. Dairy, and the number of features not in the functional category of interest within the alternative group), and performed a Hypergeometric test in R to determine functional enrichment. We adjusted for multiple testing using an FDR correction.

## Software and Data Availability

The metagenomic sequence data and associated clinical/demographic metadata were deposited in NCBI’s Sequence Read Archive under project accession number PRJNA884688. The batch and drift corrected metabolomic (and metagenomic) data is available on Github at https://github.com/krishnanlab/multiomics-embedding/blob/main/data/raw/unfiltered_micro_metab.csv. The seven embedding spaces used here, and code used to filter and merge the data, perform differential abundance and enrichment analyses, build embedding spaces, and classify microbiome and metabolite features is also available on Github at https://github.com/krishnanlab/multiomics-embedding.

The final evaluation of all generated embedding spaces is available on wandb.ai at https://wandb.ai/keenan-manpearl/multiomics_embedding?nw=nwuserkeenanmanpearl.

## Acknowledgements

We thank Diana Ir for technical assistance in shotgun metagenomic sequence analysis and all study participants in the clinical trial.

